# Power calculator for instrumental variable analysis in pharmacoepidemiology

**DOI:** 10.1101/084541

**Authors:** Venexia M Walker, Neil M Davies, Frank Windmeijer, Stephen Burgess, Richard M Martin

## Abstract

**Background:** Instrumental variable analysis, for example with physicians’ prescribing preferences as an instrument for medications issued in primary care, is an increasingly popular method in the field of pharmacoepidemiology. Existing power calculators for studies using instrumental variable analysis, such as Mendelian randomisation power calculators, do not allow for the structure of research questions in this field. This is because the analysis in pharmacoepidemiology will typically have stronger instruments and detect larger causal effects than in other fields. Consequently, there is a need for dedicated power calculators for pharmacoepidemiological research.

**Methods and results:** The formula for calculating the power of a study using instrumental variable analysis in the context of pharmacoepidemiology is derived before being validated by a simulation study. The formula is applicable for studies using a single binary instrument to analyse the causal effect of a binary exposure on a continuous outcome. A web application is provided for the implementation of the formula by others.

**Conclusions:** The statistical power of instrumental variable analysis in pharmacoepidemiological studies to detect a clinically meaningful treatment effect is an important consideration. Research questions in this field have distinct structures that must be accounted for when calculating power.

**FUNDING STATEMENT:** This work was supported by the Perros Trust and the Integrative Epidemiology Unit. The Integrative Epidemiology Unit is supported by the Medical Research Council and the University of Bristol [grant number MC_UU_12013/9]. Stephen Burgess is supported by a post-doctoral fellowship from the Wellcome Trust [100114].

**Key Messages:**

- Research questions using instrumental variable analysis in pharmacoepidemiology have distinct structures that have previously not been catered for by instrumental variable analysis power calculators.
- Power can be calculated for studies using a single binary instrument to analyse the causal effect of a binary exposure on a continuous outcome in the context of pharmacoepidemiology using the presented formula and online power calculator.
- The use of this power calculator will allow investigators to determine whether a pharmacoepidemiology study is likely to detect clinically meaningful treatment effects prior to the study’s commencement.

## INTRODUCTION

Pharmacoepidemiological studies risk irrelevance if they are insufficiently powered to detect clinically meaningful treatment effects. Prior to starting a study, the statistical power to calculate a given treatment effect can be calculated. This type of calculation is becoming increasingly important for grant and data request applications, which look to value the contribution of such studies.

The number of pharmacoepidemiology studies using instrumental variable analysis, for example with physicians’ prescribing preferences as an instrument for exposure, continues to grow.(1–6) This is partly because instrumental variable analyses have the potential to overcome some of the issues associated with conventional statistical approaches, such as residual confounding and reverse causation. As the demand to provide power calculations to support applications increases, there is a more pressing need to be able to provide power calculations for this method.

There are power calculators for instrumental variable analysis in other settings, such as Mendelian randomisation, which uses germline genetic variants as proxies for exposures in disease-related research.(7,8) However, pharmacoepidemiological research questions have distinct structures that are not sufficiently catered for by these existing calculators. Unlike Mendelian randomization studies, which often use a case-control study design, pharmacoepidemiology studies typically use a cohort study design. Further to this, pharmacoepidemiology studies usually report a risk difference for a binary exposure using a binary instrument, while Mendelian randomization studies report on a continuous exposure using a discrete or continuous genetic instrument (count of alleles or allele score respectively). As a result of these differences, as well as the stronger instruments and larger causal effects seen in pharmacoepidemiology, there is a need for a dedicated power calculator for instrumental variable analysis in the context of this field.

This paper will address how to conduct power calculations for pharmacoepidemiological studies using a single binary instrument to analyse the causal effect of a binary exposure on a continuous outcome. The formula to calculate power will be derived and then validated by a simulation study. A web application is provided for the implementation of the formula by others.

## METHODS AND RESULTS

Let us consider physicians’ prescribing preferences for two different treatments - for example a treatment of interest and a control treatment – as an instrument for exposure to these treatments. Physicians’ preferences are generally not directly observable so each physician’s prescriptions to previous patients are used as a proxy for their preferences. This results in a binary instrument that takes a value of one if the physician issued a prescription for the treatment of interest to their previous patient and a value of zero if they prescribed the control treatment. We will derive the formula for the power of studies that use this instrument to measure the causal effect of a drug exposure on a continuous outcome, for example systolic blood pressure or low density lipoprotein cholesterol.

### FORMULA DERIVATION

The instrumental variable analysis we consider requires the following three variables; namely a binary instrument ***Z***, a binary exposure ***X*** and a continuous outcome ***Y***. The outcome for patient ***i***, for ***i* = 1, …**, ***n***, is modelled as follows:

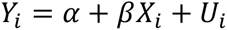

where ***U_i_*** is a zero-mean error term containing unobserved confounders, determining both the outcome ***Y_i_*** and the treatment ***X_i_***. The instrument ***Z_i_*** affects treatment ***X_i_***, but is not associated with the unobserved confounders and has no direct effect on the outcome.

Let 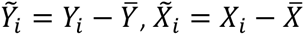 and 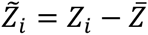, where 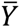, 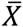 and 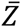 are sample averages. Denote by 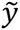, 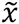 and 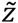 the ***n***-vectors of observations on 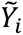, 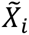 and 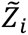 respectively. The two-stage least squares (2SLS) estimator of ***β*** is then given by:

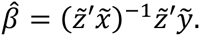

The variance of the 2SLS estimator is:

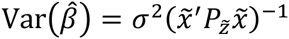

where 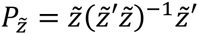 and 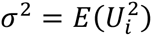 is the residual variance. Note that conditional homoscedasticity holds so the variance is constant for all values of the instrument i.e. 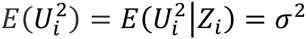. for ***i* = 1,…, *n***.

Consider the term 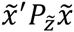:

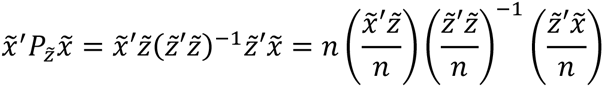

Let ***p_Z_* = *P***(***Z* = 1)**, ***p_X_* = *P***(***X* = 1)** and ***p_XZ_* = *P***(***X* = 1|*Z* = 1)**. In large samples:

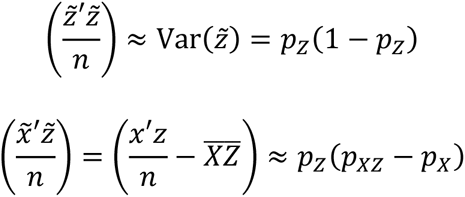

Hence 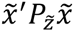 can be presented in the following way:

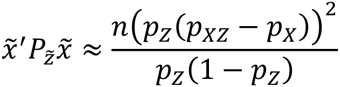

Now consider the instrumental variable estimator of ***β***. Using the asymptotic distribution 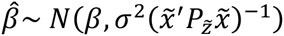, the distribution of the t-test statistic under the null hypothesis ***H*_0_: *β* = *β*_0_** is:

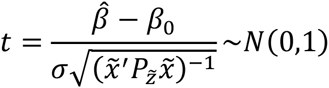

The distribution of the test statistic under the alternative hypothesis ***H*_1_: *β* = *β*_0_ + *δ*** is:

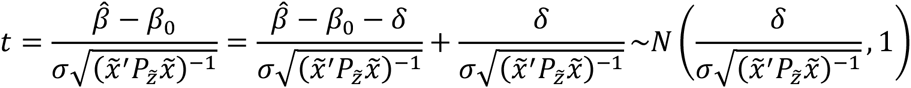

The null hypothesis is rejected if **|*t*| > *c*_*α*_** where ***c***_***α***_ is the critical value at significance level ***α***.

The power is the probability the test statistic will exceed the critical value, which is:

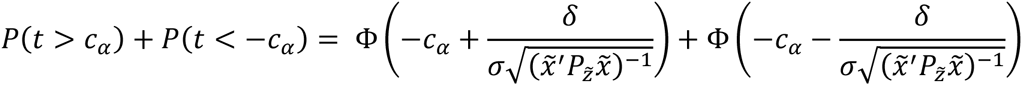

where **Φ(*s***) is the cumulative standard normal distribution function evaluated at ***s***.

Power therefore increases as the value of ***σ*** decreases and/or the value of 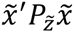 increases. By substituting 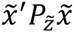 and simplifying, we obtain the following formula for power:

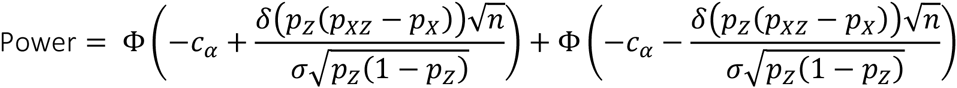

The formula requires a total of seven parameters to be specified. These are the significance level ***α***, the size of the causal effect ***δ***, the residual variance 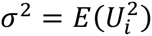, the frequency of the instrument ***p_Z_* = *P***(***Z* = 1)**, the frequency of exposure ***p_X_* = *P***(***X* = 1)**, the probability of exposure given the instrument ***p_XZ_* = *P***(***X* = 1|*Z* = 1)** and the sample size ***n***. Note that the following must hold:

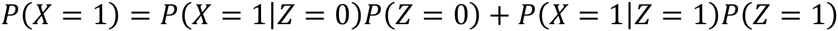

This formula for power is available for use via an online calculator, which can be found at https://venexia.shinyapps.io/PharmIV/.

Note that the frequency of exposure in an instrumental variable analysis of this type is likely to be higher than a general population study because a drug is compared against one or more other drugs in a population of people with the indication for these treatments. General population studies on the other hand tend to compare a population who received the drug of interest with a population who did not receive it and consequently the frequency of exposure is generally much lower. The effect of varying the parameters within the formula on a study’s power is best presented graphically. figure 1 illustrates an example of the effect of the frequency of the exposure ***p_X_* = *P***(***X* = 1)** on the power of a study to detect a causal effect of **δ = –0.15** using an instrument with a frequency of ***p_Z_* = 0.20**, a residual variance of ***σ*^2^ = 1** and a sample size of up to 30,000 participants. Both increasing the frequency of exposure up to 50% and increasing the sample size results in increased power for this study.

**figure 1:**
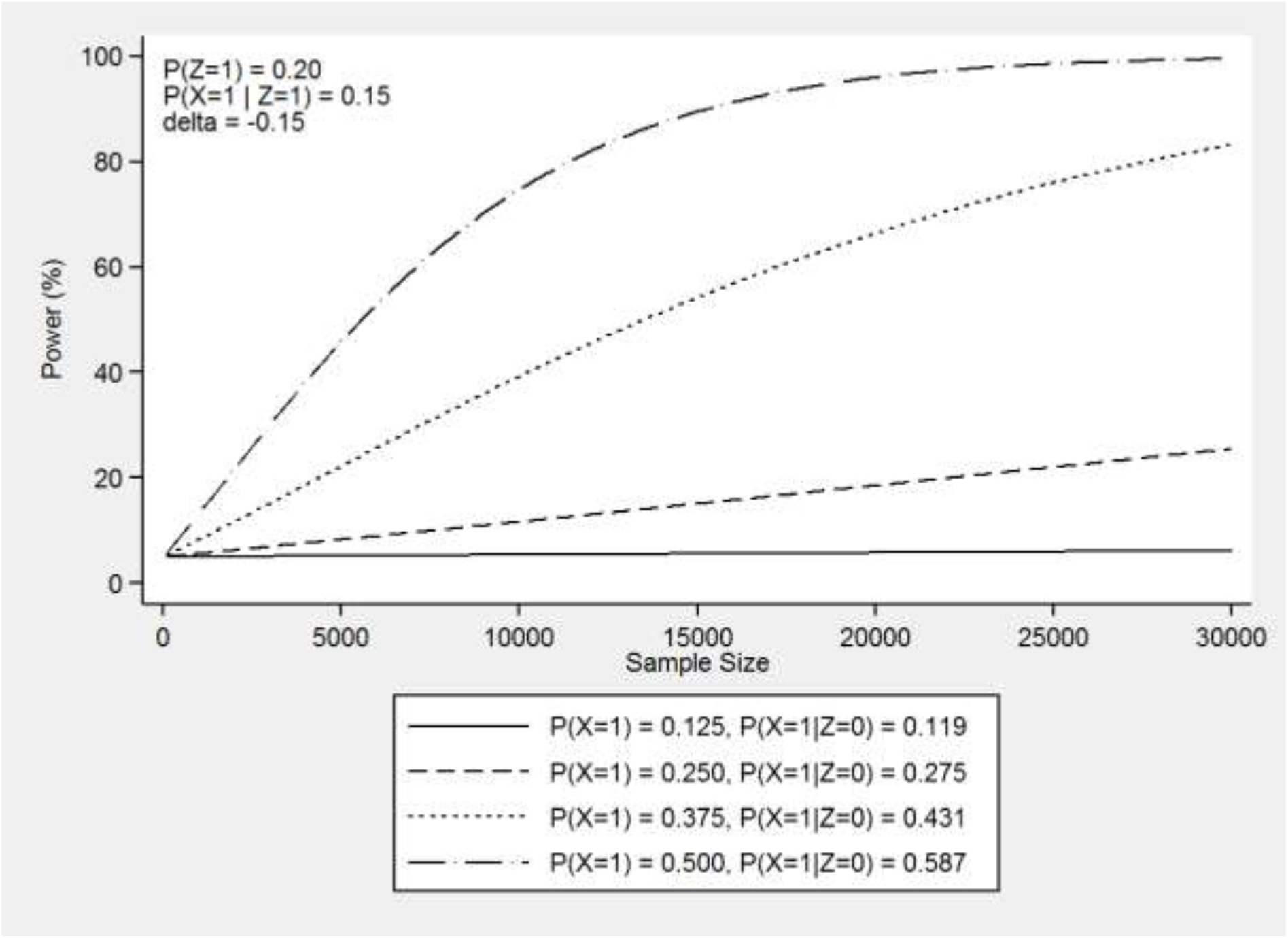
Power curves for several values of the frequency of exposure ***p_X_* = *P***(***X* = 1)** that show the effect on the power of a study to detect a causal effect of ***δ* = –0.15** using an instrument with a frequency of ***p_Z_* = 0.20**, a residual variance of ***σ*^2^ = 1** and a sample size of up to 30,000 participants.

### FORMULA VALIDATION

To validate the power formula, we conducted a simulation. We simulated the data by defining the three variables necessary to conduct instrumental variable analysis with a single instrumental variable as follows:

Instrument: ***Z_i_*** ∼ Binomial (1, ***p_Z_***)

Exposure: 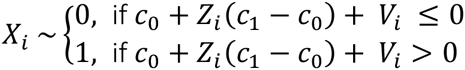

Outcome: ***Y_i_*** ∼ ***δX_i_*** + ***U_i_***

Where ***p_Z_* = *P***(***z* = 1)** is the frequency of the instrument, **c_j_ = Φ^−1^P(*X* = 1|*Z* = *j*))** for ***j* = 0,1** are the inverse cumulative standard normal distribution, or quantile, functions of the conditional probabilities of exposure given the instrument, ***δ*** is the causal effect, and ***U_i_*** and ***V_i_*** are standard normally distributed error terms with covariance ***ρ***.

The formula uses a binary instrument, binary exposure and continuous outcome and so the above variables were simulated to recreate data of this form. The instrument ***Z*** is modelled by a binomial distribution parameterised by its frequency ***p_Z_* = *P***(***Z* = 1)**. This ensures a binary variable with the correct probability of success. The exposure ***X*** is also binary but is modelled using a threshold model. The variability in the equation for the exposure comes from the normally distributed error term ***V_i_***. The use of the model equation allows the exposure ***X*** to be associated with the instrument ***Z***. The outcome ***Y*** is modelled by its model equation ***Y_i_* = *δX_i_*** + ***U_i_***. In the model, the instrument is valid as the outcome ***Y*** is only associated with the exposure ***X***, as dictated by the causal effect ***δ***, and is not associated with the instrument ***Z*** other than through the exposure ***X***.

Using the generated data, we performed an instrumental variable analysis using the command IVREG2 in Stata.(9) From this analysis, we recorded the coefficient of the exposure ***X*** with the 95% confidence interval. We then counted the number of simulations for which the confidence interval excluded the null and divided this by the total number of simulations to determine the power. By running the simulation and calculating the formula using the same parameters, we are able to validate the formula against the simulation.

We present the power calculated from both the simulation and the formula for several parameter combinations in Table 1. The table contains 27 different simulations and each was repeated 10,000 times. The simulations consider each combination of three values of the frequency of exposure ***p_X_* = 0.10, 0.25, 0.50**, three values of the probability of exposure given the instrument ***p_XZ_* = 0.15, 0.30, 0.45**, and three values of the sample size ***N* = 10000, 20000, 30000**. We set the frequency of the instrument ***p_Z_* = 0.20**, the causal effect ***δ* = –0.15**, the residual variance ***σ*^2^ = 1** and calculated ***P***(***X* = 1|*Z* = 0)** according to the following equation:

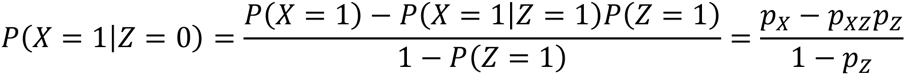

The Stata code used to create the simulation is available online at https://github.com/venexia/PharmIV. The effect of confounding was removed as a parameter because the power was insensitive to its value in the simulation setting. Details of the simulations conducted to test this can be found in Supplementary File 1.

### SIMULATION RESULTS

**Table 1:** A comparison of the power calculated from the formula and a validation simulation for an instrumental variable analysis where the causal effect ***δ* = –0.15**, the frequency of the instrument ***p_z_* = 0.20** and the residual variance ***σ*^2^ = 1**.

| $p_X$ | $p_{XZ}$ | 10,000 patients | | 20,000 patients | | 30,000 patients | |
| --- | --- | --- | --- | --- | --- | --- | --- |
|  |  | Formula | Simulation | Formula | Simulation | Formula | Simulation |
| 0.10 | 0.15 | 6.6% | 6.1% | 8.3% | 7.9% | 10.0% | 9.8% |
|  | 0.30 | 32.3% | 33.3% | 56.4% | 55.5% | 73.8% | 73.9% |
|  | 0.45 | 74.7% | 75.6% | 96.0% | 95.9% | 99.5% | 99.5% |
| 0.25 | 0.15 | 11.7% | 11.4% | 18.6% | 18.3% | 25.5% | 25.5% |
|  | 0.30 | 6.6% | 5.4% | 8.3% | 7.9% | 10.0% | 9.8% |
|  | 0.45 | 32.3% | 32.8% | 56.4% | 56.1% | 73.8% | 73.7% |
| 0.50 | 0.15 | 74.7% | 74.2% | 96.0% | 95.9% | 99.5% | 99.6% |
|  | 0.30 | 32.3% | 32.5% | 56.4% | 57.1% | 73.8% | 73.7% |
|  | 0.45 | 6.6% | 5.0% | 8.3% | 7.2% | 10.0% | 10.1% |

The formula and the simulation consistently provide similar results with an absolute mean difference of **0.4%** for the parameter combinations presented in Table 1. There is also no discernible pattern in the differences suggesting systematic bias is not present. Further to this, the power is consistent with its behaviour in other established power calculations. For example, increasing sample size universally improves power for all parameter combinations.

## DISCUSSION

In this paper, we have derived the formula necessary to calculate power for instrumental variable analysis with a single binary instrument, binary exposure and continuous outcome in the context of pharmacoepidemiology. The formula has been shown to be valid by comparison against a simulation study, which concluded that the formula provided near true values across a range of realistic parameters.

As for any power formula, the formula presented here is limited by its parameters, which simplify the dataset being considered. Power calculated from such formulae cannot account for dataset characteristics outside of these parameters. For example, the formula makes no allowance for the presence of missing data – a known limiting factor on the power of a study. By allowing for missing data in the anticipated sample size, conservative estimates for the power of a study can be obtained using the formula presented. Further work is needed in order to establish the formula for power in other scenarios that use instrumental variable analysis within a pharmacoepidemiology context. This includes analyses with binary outcomes and analyses that involve multiple instrumental variables.

As the use of instrumental variable analysis in pharmacoepidemiology becomes more commonplace, there is an increasing need to provide power calculations for studies using this type of analysis. To provide such information, accessible and accurate power formulae need to be made available. By using the formula presented here and the online tool, it is hoped that pharmacoepidemiologists can calculate the power of instrumental variable analysis studies with a single binary instrument, binary exposure and continuous outcome with ease.

